## Supplementary material for "CoSMIC - A hybrid approach for large-scale, high-resolution microbial profiling of novel niches": Supp CoSMIC

### Supplementary data

Code is available at https://github.com/rezenman/CoSMIC
Raw reads are available at
https://www.dropbox.com/sh/kyaecrfnpkdxd96/AACANythrtzEBQApCalFzwUza?dl=0

#### Figures S1-5

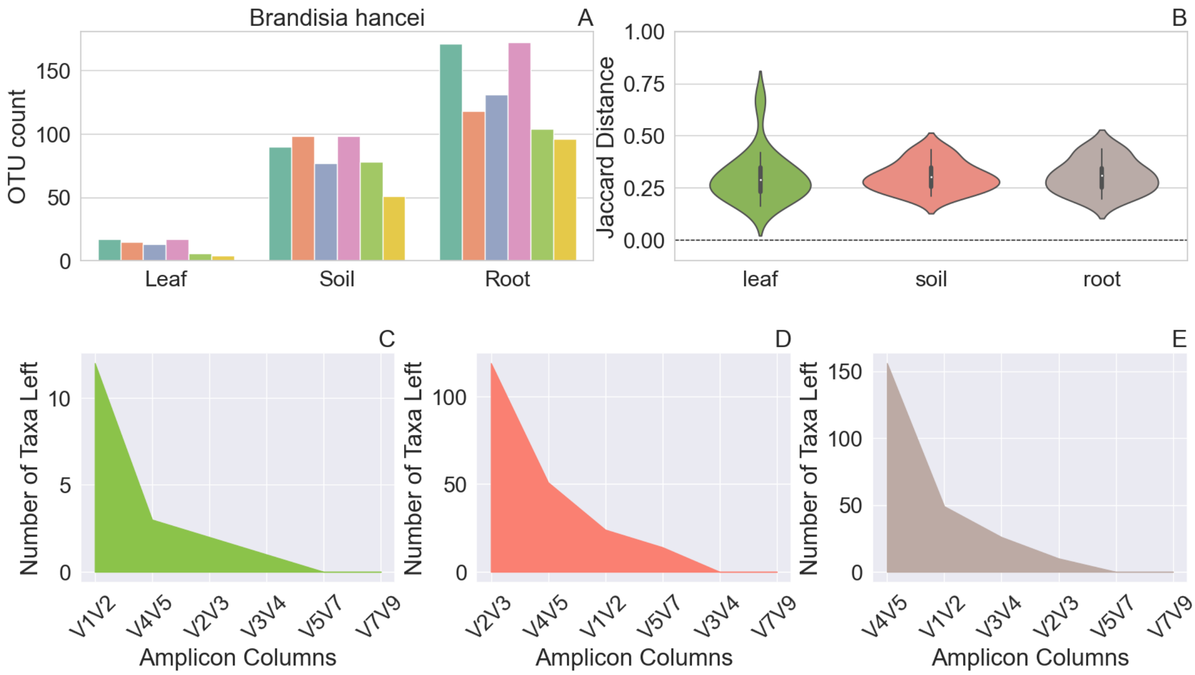


**Figure S1. Microbial profiling of *Brandisia hancei* depends on the applied amplicon.** Leaf, soil, and root samples were profiled by the Qiagen kit that provides six amplicons along the 16S rRNA gene and subsequently analyzes their reads using the CLC Genomics platform. **A.** The number of OTUs detected for soil, leaf, and root samples of each plant for each of the six amplicons. **B.** Distribution of Jaccard distances among pairs of amplicons. In each pair, e.g., V1V2 vs. V3V4, a Jaccard distance is calculated between the lists of taxons detected by CLC Genomics in each region. **C-E.** Decay graph of the number of unique taxons identified by the amplicons analyzed. Amplicons are ordered by the number of unique taxons a region contributes to the former set of regions. C-Leaf, D-Soil, and E-Root.

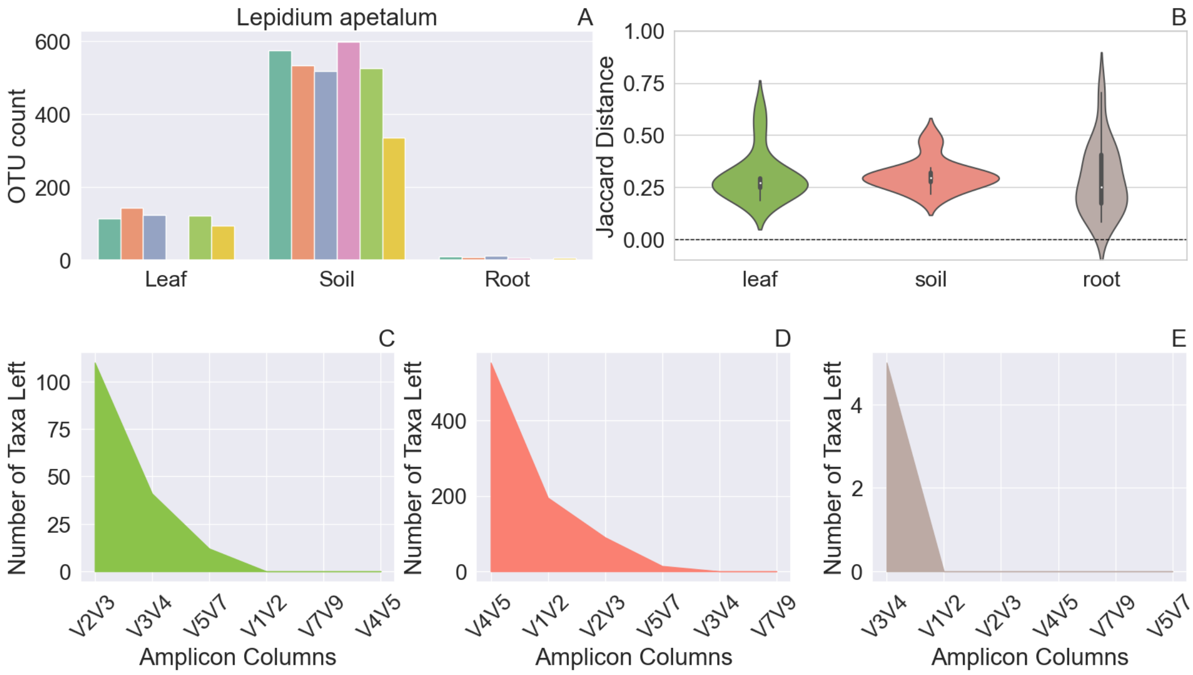


**Figure S2. Analogous figure to Fig. S1, for *Lepidium apetalum***

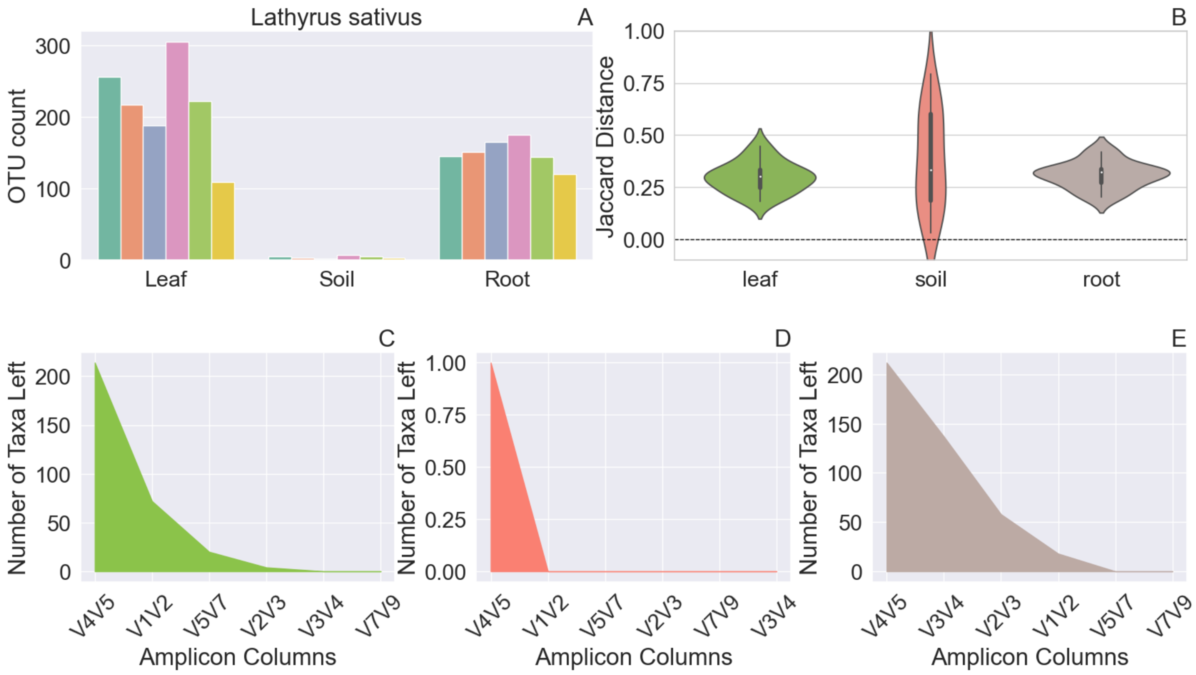


**Figure S3. Analogous figure to Fig. S1, for *Lathyrus sativus***

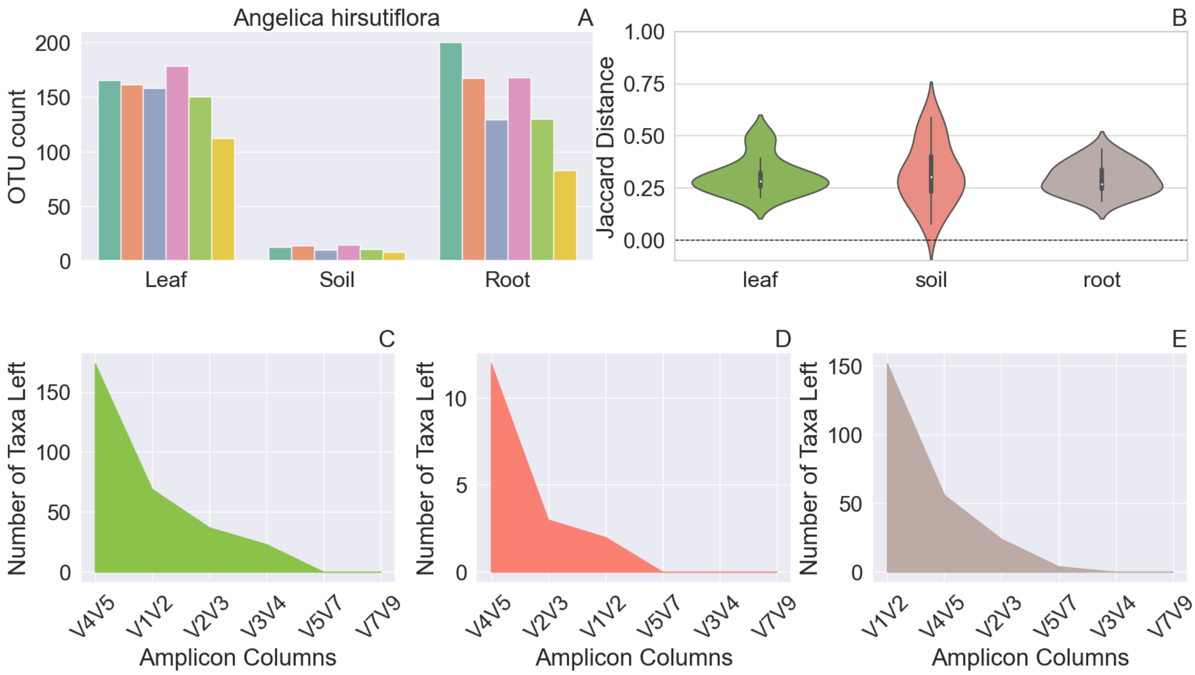


**Figure S4. Analogous figure to Fig. S1, for *Angelica hirsutiflora***

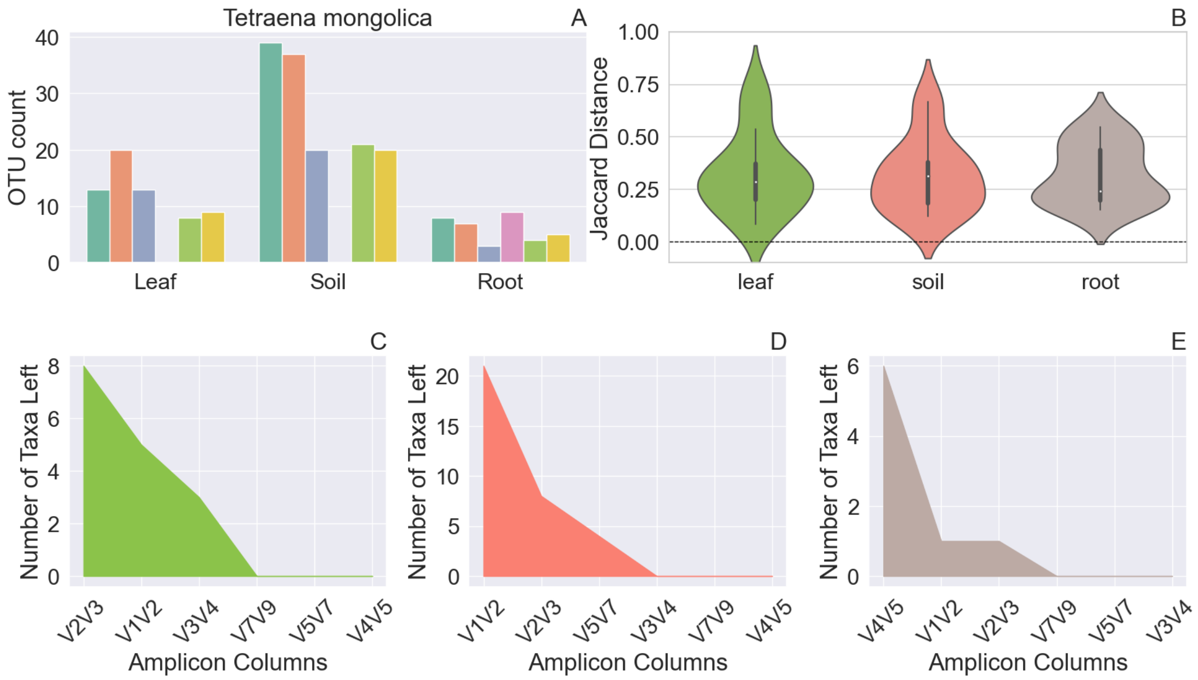


**Figure S5. Analogous figure to Fig. S1, for *Tetraena mongolica***

#### Figures S6-8

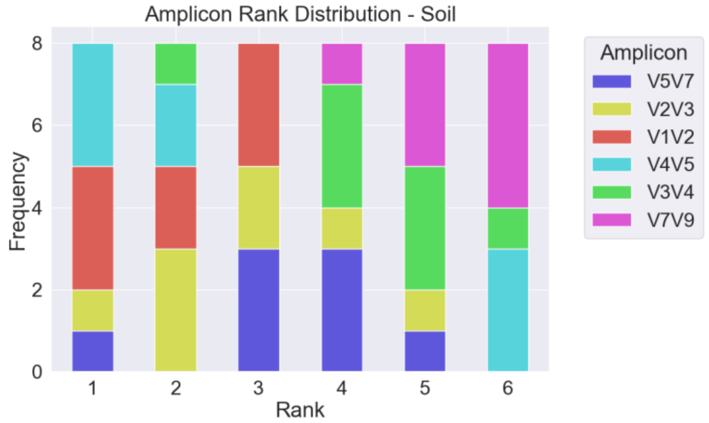


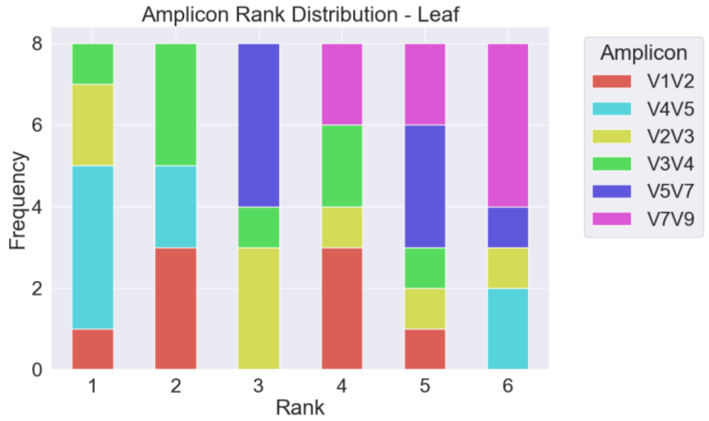


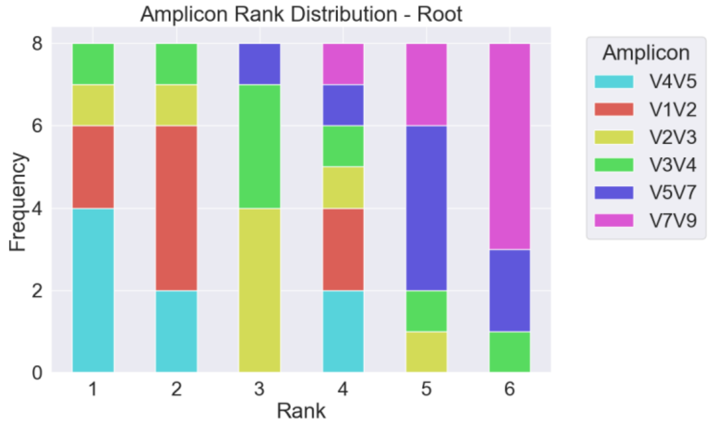


**Figures S6-8. Ranking of Amplicons used in Qiagen analysis.** Analysis of leaf, soil, and root samples involved six distinct amplicons. In this figure, each amplicon is ranked according to the amount of information it contributed to the analysis. There is no evident ranking, thus indicating that the information contribution for each measurement originates from varying amplicons.

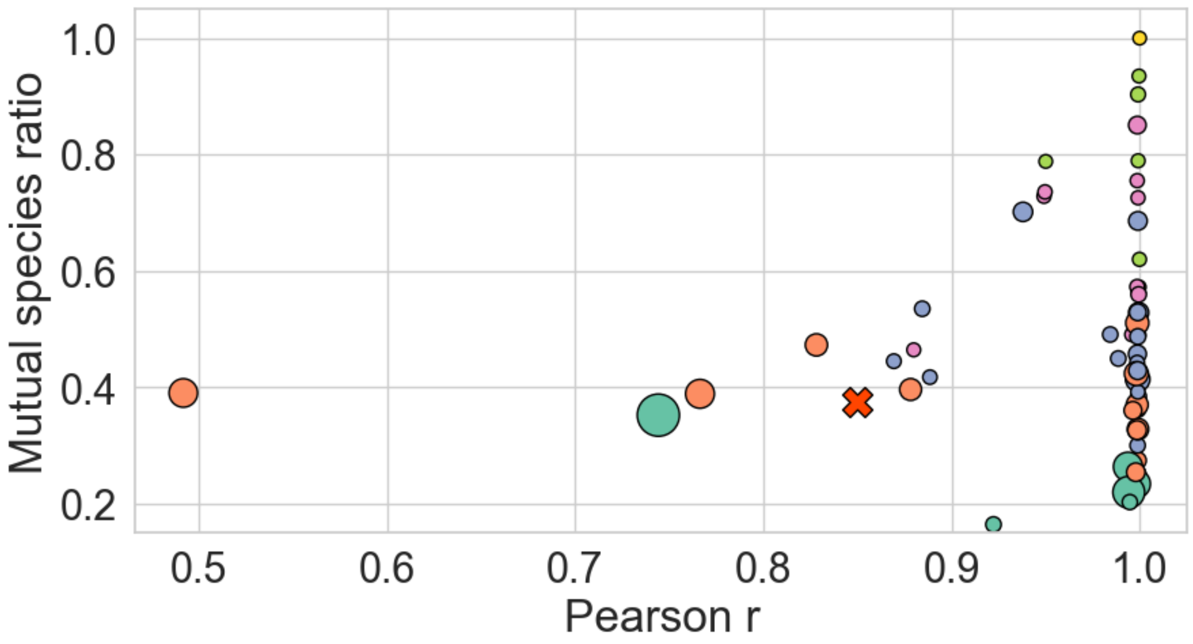


**Figure S9. A comparison of amplicon sets for leaf samples.** Each dot corresponds to a specific region combination averaged across samples, where the marker size scales with the ambiguity and axes correspond to Pearson correlation (horizontal) and mutual species (vertical). A marker's color indicates the number of amplicons used for SMURF analysis. Hence, a single marker corresponds to 6 regions (yellow), and 6 green dots correspond to the different groups of 5 regions. The marker "x" corresponds to a specific set of 3 regions comprising V1V2, V2V3, and V4V5.

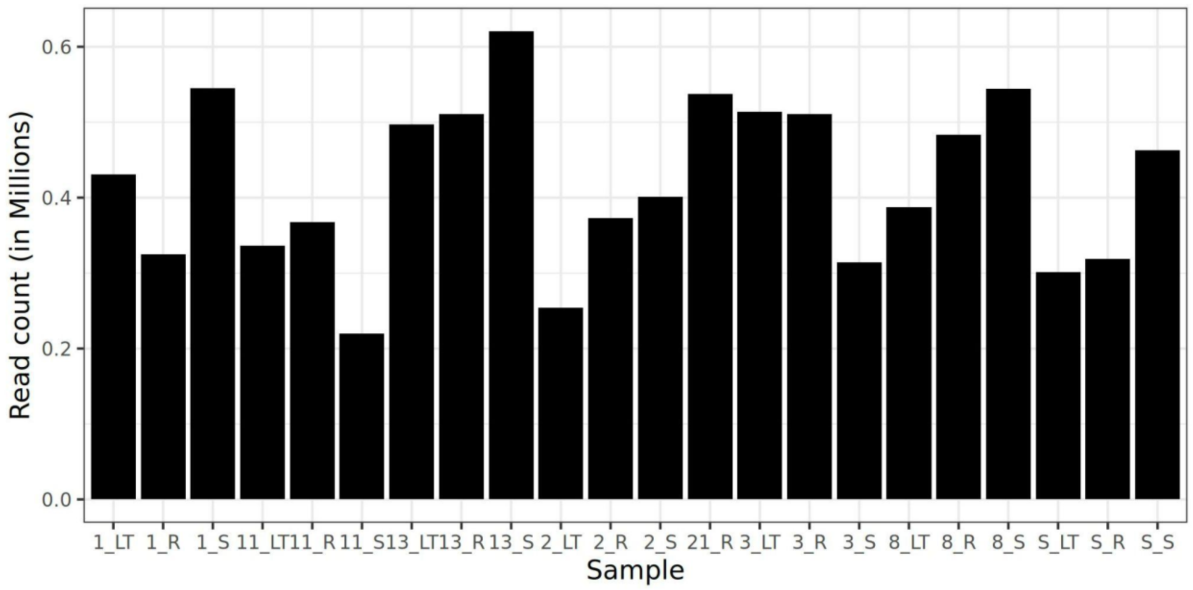


**Figure S10. Number of Illumina reads for Qiagen QIAseq sequenced samples.** Number of raw Illumina reads for each sample prepared using the Qiagen QIA seq kit, which amplifies 6 variable regions along the 16S rRNA gene. Samples' descriptions appear in Table S1.

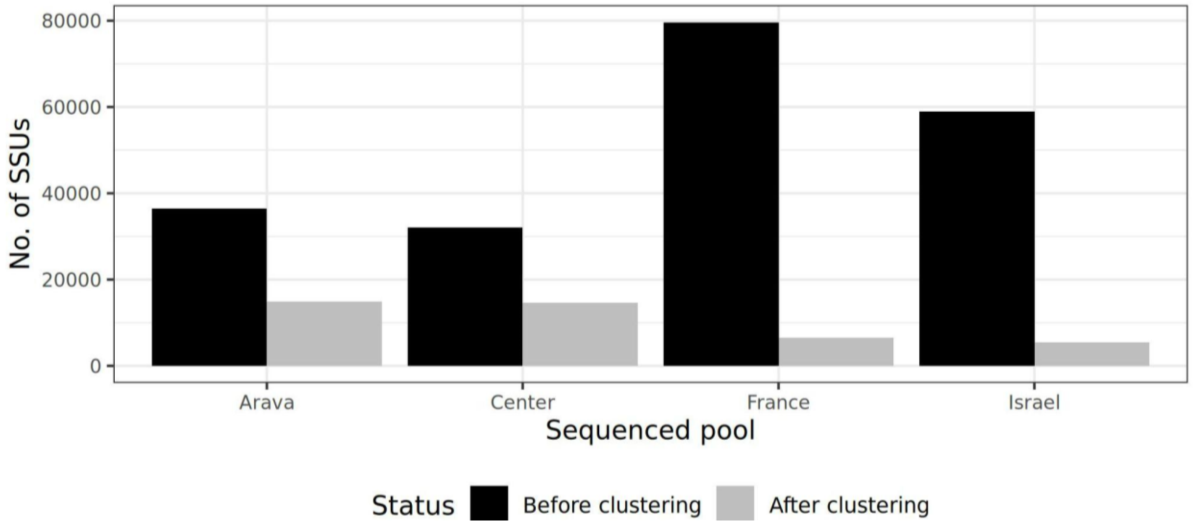


**Figure S11. Number of SSU derived from PacBio long read sequencing.** In order to reduce redundancy of amplicons derived from the PacBio long read sequencing we clustered each sample using cd-hit with an identity threshold of 99%. In black is the number of sequences derived for each sample prior clustering; in gray is the number of seeds obtained after clustering.

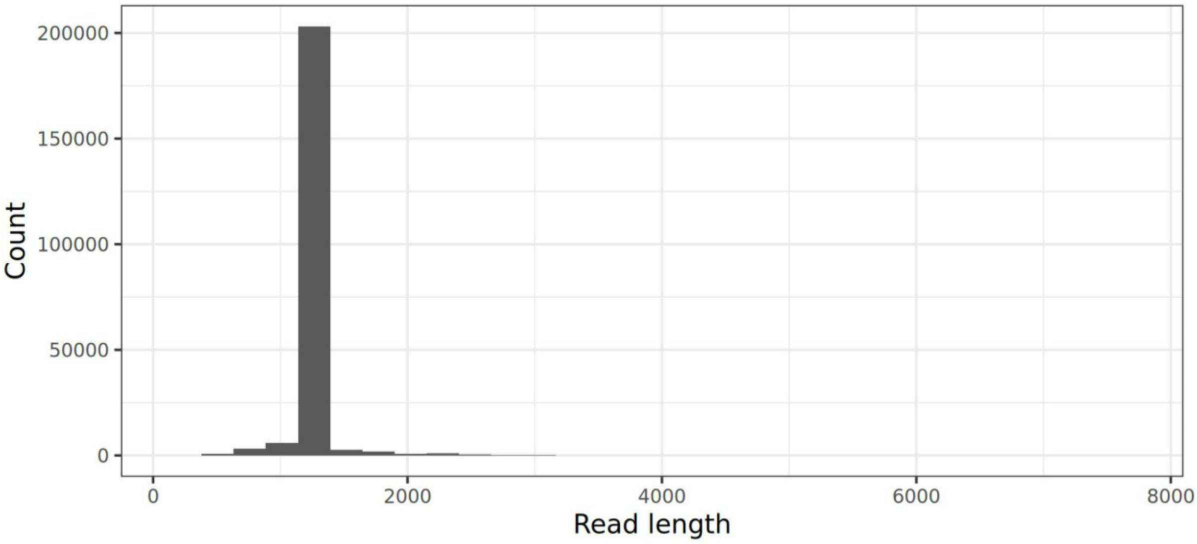


**Figure S12. Read length distribution for PacBIo long reads sequences.** Read length distribution of LNA-based 16S rRNA amplicons, obtained from the LNA amplification of all four pools of samples (*Spongia officinalis* from France and Israel; and plants from Israel's lowland and Arava regions).

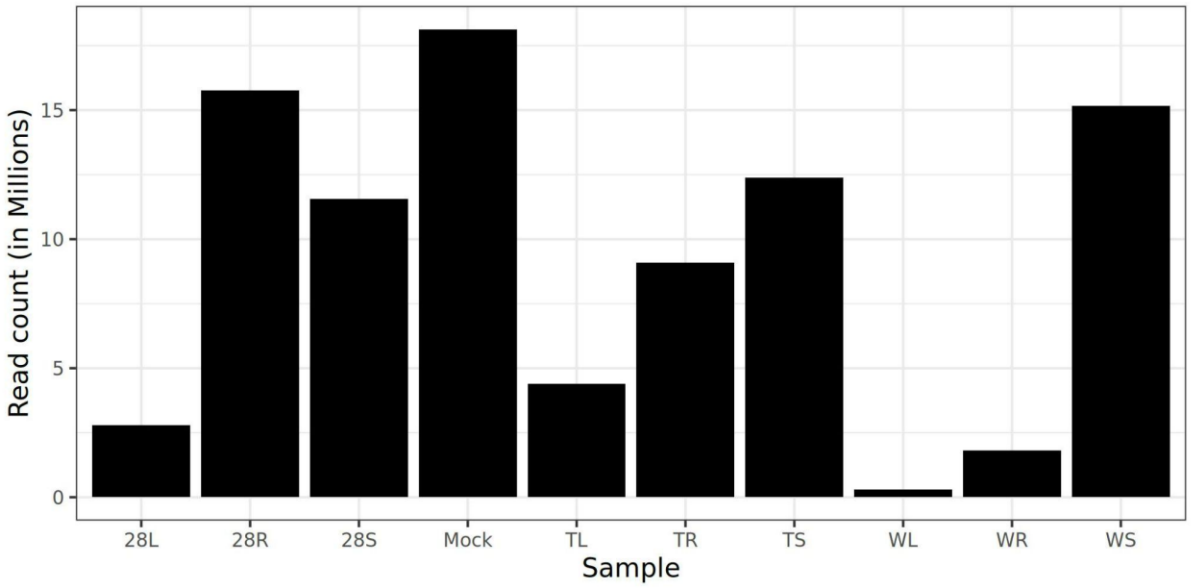


**Figure S13. Number of reads for samples sequenced using Loop Genomics synthetic long reads.**

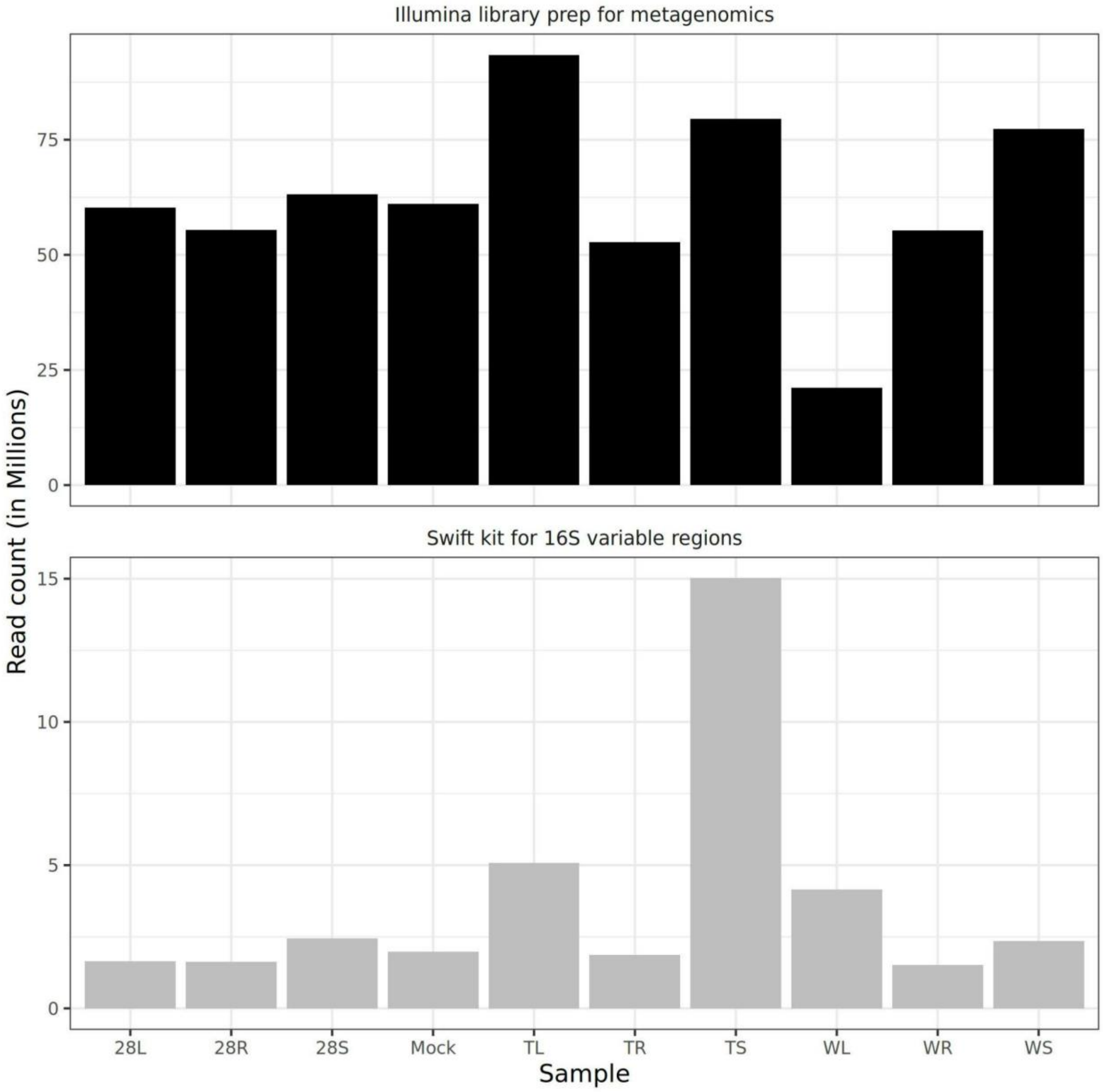


**Figure S14. Number of Illumina reads obtained for each sample.** For each sample (horizontal), the number of raw reads obtained by illumina short read sequencing is presented on the vertical axis (in millions). The upper panel corresponds to samples prepared for metagenomics analysis while the lower panel corresponds to samples prepared using the Swift Biosciences kit, which amplifies multiple regions of the 16S rRNA gene.

**Table S1. List of samples used in the study.** *Organism* refers to the organism from which the sample was taken; *Tissue,* refers to the tissue sampled in the organism; *Sample ID*, refers to sample name as recorded in our data; *Sample preparation kit*, refers to the commercial kit used to prepare the samples for microbiome analysis; *Relevant results section*, refers to the part in the results where we used the listed samples; *Description*, refers to whether this entry is a single sample or pool of samples. Asterix refers to pools of multiple samples as depicted in the methods section and not individual samples.

| **Organism** | **Tissue** | **Sample ID** | **Sample preparation kit** | **Relevant results section** | **Description** |
|---|---|---|---|---|---|
| *Lepidium apetalum* | Soil | 1_S | Qiagen QIAseq | Combining results from several variable regions partially overcomes problems in profiling | Single sample |
| | Leaf | 1_LT | Qiagen QIAseq | | Single sample |
| | Root | 1_R | Qiagen QIAseq | | Single sample |
| *Lepidium apetalum* | Soil | 2_S | Qiagen QIAseq | | Single sample |
| | Leaf | 2_LT_5 | Qiagen QIAseq | | Single sample |
| | Leaf | 2_LT_8 | Qiagen QIAseq | | Single sample |
| | Root | 2_R | Qiagen QIAseq | | Single sample |
| *Aster poliothamnus* | Soil | 3_S | Qiagen QIAseq | | Single sample |
| | Leaf | 3_LT | Qiagen QIAseq | | Single sample |
| | Root | 3_R | Qiagen QIAseq | | Single sample |
| *Brandisia hancei* | Soil | 8_S_1 | Qiagen QIAseq | | Single sample |
| | Soil | 8_S_4 | Qiagen QIAseq | | Single sample |
| | Leaf | 8_LT | Qiagen QIAseq | | Single sample |
| | Root | 8_R | Qiagen QIAseq | | Single sample |

| | | | | | |
|---|---|---|---|---|---|
| *Tetraena mongolica* | Soil | 11_S | Qiagen QIAseq | | Single sample |
| | Leaf | 11_LT | Qiagen QIAseq | | Single sample |
| | Root | 11_R | Qiagen QIAseq | | Single sample |
| *Angelica hirsutiflora* | Soil | 13_S | Qiagen QIAseq | | Single sample |
| | Leaf | 13_LT | Qiagen QIAseq | | Single sample |
| | Root | 13_R | Qiagen QIAseq | | Single sample |
| | Root | 21_R | Qiagen QIAseq | | Single sample |
| *Lathyrus sativus* | Soil | S_S | Qiagen QIAseq | | Single sample |
| | Leaf | S_LT | Qiagen QIAseq | | Single sample |
| | Root | S_R | Qiagen QIAseq | | Single sample |
| *Hymelaea hirsuta* | Soil | 28S | Swift 16S+ITS PANEL | Experimental evaluation of CoSMIC | Single sample |
| | Leaf | 28L | Swift 16S+ITS PANEL | | Single sample |
| | Root | 28R | Swift 16S+ITS PANEL | | Single sample |
| *Solanum lycopersicum* | Soil | TS | Swift 16S+ITS PANEL | | Single sample |
| | Leaf | TL | Swift 16S+ITS PANEL | | Single sample |
| | Root | TR | Swift 16S+ITS PANEL | | Single sample |
| *Triticum aestivum* | Soil | WS | Swift 16S+ITS PANEL | | Single sample |
| | Leaf | WL | Swift 16S+ITS PANEL | | Single sample |
| | Root | WR | Swift 16S+ITS PANEL | | Single sample |

| | | | | | |
|---|---|---|---|---|---|
| Mock community | Mock | Mock | Swift 16S+ITS PANEL | | Mock community built from 12 known species (listed in the methods section) |
| *Spongia officinalis* - France* | | France | PacBio SMRTbell Express template Prep Kit 2.0 | Enriching the database using long-read sequencing | Pooled samples from the Mediterranean sea in Marseille, France* |
| *Spongia officinalis* - Israel* | | Israel | PacBio SMRTbell Express template Prep Kit 2.0 | | Pooled samples from the Mediterranean sea near Haifa, Israel* |
| Arava* | | Arava | PacBio SMRTbell Express template Prep Kit 2.0 | | Pooled plant samples from the desert region* |
| Lowlands pool* | | Center | PacBio SMRTbell Express template Prep Kit 2.0 | | pooled plant samples from the lowlands region* |
| Loop pool* | | Loop_LNA | Loop genomics LoopSeq PCR Amplicon Kit | | Pooled plant samples used in Qiagen and Swift Analysis* |
